# Chromosome evolution at the origin of the ancestral vertebrate genome

**DOI:** 10.1101/253104

**Authors:** Christine Sacerdot, Alexandra Louis, Céline Bon, Hugues Roest Crollius

## Abstract

About 450 million years ago, a marine chordate was subject to two successive whole genome duplications (WGDs) before becoming the common ancestor of vertebrates and diversifying into the more than 60,000 species found today. Here, we reconstruct in details the evolution of chromosomes of this early vertebrate along successive steps of the two WGD. We first compared 61 extant animal genomes to build a highly contiguous order of genes in a 326 million years old ancestral *Amniota* genome. In this genome, we established a well-supported list of duplicated genes originating from the WGDs to link chromosomes in tetrads, a telltale signature of these events. This enabled us to reconstruct a scenario where a pre-vertebrate genome composed of 17 chromosomes duplicated into 34 chromosomes, and was subject to 7 chromosome fusions before duplicating again into 54 chromosomes. After the separation of *Agnatha* (jawless fish) and *Gnathostomata*, four more fusions took place to form the ancestral *Euteleostomi* genome of 50 chromosomes. These results firmly establish the occurrence of the two WGD, resolving in particular the ambiguity raised by the analysis of the lamprey genetic map. In addition, we provide insight into the origin of homologous micro-chromosomes found in the chicken and the gar genomes. This work provides a foundation for studying the evolution of vertebrate chromosomes from the standpoint of a common ancestor, and particularly the pattern of duplicate gene retention and loss that resulted in the gene composition of extant genomes.

## INTRODUCTION

New gene copies appear by small scale duplication during genome evolution (Innan and Kondrashov 2010), contributing to genetic innovation and phenotypic diversity (Conrad 2008). But whole genome duplications (WGDs) are rarer and more dramatic event, especially in vertebrate evolution, in contrast to plants where they appear to be more frequent (Van de Peer et al. 2009). Yet about 450 million years ago, prior to its diversification into more than 60,000 extant species, an early vertebrate lineage was subject to two WGD in relatively rapid succession. Envisioned by Susumo Ohno since the early 70’s (Ohno 1970), these events known as the “2R scenario” have since been firmly established by at least three genome wide studies. The first demonstration of the 2R showed that a large part of the human genome was covered by a clear pattern of four-way paralogous regions, defined by a subset of paralogous genes that were duplicated before the fish-tetrapod split (Dehal and Boore 2005). Soon after, a first version of the ancestral vertebrate pre-2R proto-chromosomes was reconstructed using duplicated regions of the human genome, that contains four copies of the ancestral vertebrate genome (Nakatani et al. 2007). This study proposed a scenario starting from 10-13 protochromosomes, each duplicating twice into 40 chromosomes during the 2R, but including three chromosome fissions and the loss of six chromosomes. Finally, the comparison of the cephalochordate amphioxus genome (*Branchiostoma floridae*) with the genome of several vertebrates defined the unordered gene content of 17 linkage groups of the last common chordate ancestor, which revealed a global four-fold conserved macro-synteny with vertebrate genomes (Putnam et al. 2008).

Approximately 35% of extant human genes still exist in duplicate copies (ohnologs) owing to the 2R WGDs (Makino and McLysaght 2010; Singh et al. 2015), including the four clusters of HOX genes in tetrapods (Amores et al. 1998). Ohnologs have been shown to be enriched in disease genes (Makino and McLysaght 2010; Singh et al. 2012; McLysaght et al. 2014; Rice and McLysaght 2017) and to influence the frequency of structural variations in human populations (Makino et al. 2013). The ancient 2R genome duplications therefore still exert a strong influence on present-day genomes, warranting a better understanding of their origin. The first reconstruction of the evolution of the vertebrate karyotype during the 2R (Nakatani et al. 2007) left a number of questions unanswered. Its resolution could not determine if chromosome fusions or fissions took place between the 2 WGDs to explain a number of conserved linkage patterns with respect to extant chromosomes. Some predicted ancestral vertebrate chromosomes could not be identified in extant genomes, either because of their rapid loss or because the genome sequences that were available at the time might not have made it possible to identify all duplicated chromosomes. Indeed, more than 400 millions years have elapsed since the vertebrate ancestor (*Vertebrata*), during which many chromosome rearrangements have occurred, blurring the initial signal. In addition, a recent comparison between the lamprey genetic map and the chicken genome questioned the 2R scenario by suggesting that a single WGD and multiple segmental duplications could explain the macrosynteny patterns observed (Smith and Keinath 2015). Both the uncertainties of the first reconstruction of the pre-2R genome and this alternative to the 2R scenario motivate a detailed analysis of the early evolution of vertebrate chromosomes.

The number of sequenced vertebrate genomes has recently greatly increased, allowing the reconstruction of ancestral genomes with much higher accuracy. Here, we used the AGORA algorithm to reconstruct the ancestral karyotype (gene content, order and orientation) of *Amniota* (326 My), ancestral to birds, reptiles and mammals, to reexamine the question of early vertebrate evolution.*Amniota* is much closer to the ancestral vertebrate than extant species, and should theoretically exhibit a comparatively much clearer signature of any genomic event that took place in early vertebrate evolution. After identifying thousands of ohnologous genes in the *Amniota* genome, we applied a strategy (outlined in Supplemental Figure S1) to identify tetrads of chromosome segments, each resulting from two successive duplications. With additional information from the more ancient chordate proto-chromosomes reconstructed using the comparison of amphioxus with vertebrate genomes (Putnam et al. 2008), we dissected the events that took place between the 2 WGD. We identified seven chromosome fusions that took place in that interval, and did not identify any chromosome fissions. This result leads to the conclusion that the pre-2R vertebrate genome contained 17 chromosomes and that all vertebrates descend from a post-2R genome comprising 54 chromosomes. We show that the human genome still bears a strong imprint of the pre-2R genome, and provide resources to study the impact of the 2R in vertebrates from the perspective of a founding reference point.

## RESULTS

### Identification of pairs of ohnologous genes in the *Amniota* genome

We reconstructed the *Amniota* karyotype using AGORA (methods), starting from 19786 gene identifiers inferred from Ensembl gene trees to have existed in the *Amniota* genome. In this reconstruction, genes are ordered and oriented in scaffolds, and the current level of assembly is composed of 470 such scaffolds, with 50% larger than 253 genes (N50 length). This high level of contiguity prompted us to select the 56 scaffolds larger than 50 genes as an initial set of contiguous ancestral regions (CARs; mean CAR length 256 genes, 12134 genes total) to start identifying duplicated regions.

Genes that duplicated during the 2R WGD, called ohnologs, are key to identifying pairs of duplicated chromosome segments. Two major studies have identified ohnologs from the 2R in the human and other vertebrate genomes (Makino and McLysaght 2010; Singh et al. 2015). Both used conserved synteny and homology to identify these duplicates in extant genomes. We remapped these on the *Amniota* genome via their respective gene trees; the “MM” list was deduced from that of (Makino and McLysaght 2010) and contained 4,870 gene pairs; the three “SAI” lists were built using the ohnolog pairs of(Singh et al. 2015) with three levels of confidence as defined in the original article (strict, intermediate and relaxed) and contained respectively 2,873 (“SAI-strict”), 5,253 (“SAI-inter”) and 7,806 (“SAI-relax”) *Amniota* ohnolog pairs. These previously published lists were established using human genes, which underwent extensive evolutionary events (sequence diversification, deletions and small scale duplications) since the WGDs. To bypass some of these issues, we also identified ohnolog pairs directly in the reconstructed *Amniota* genome using phylogenetic trees and gene coordinates in their respective CARs, resulting in a “DYOGEN” list containing 5,616 pairs (see methods).

The five lists of ohnolog pairs overlap poorly (12% of the union of the 5 lists), but the corresponding lists of ohnolog genes show a better overlap (25%; Figure 1A). This is because the lists of ohnolog pairs group the same genes as different pairs, a situation that can happen because after two rounds of WGD each original gene is duplicated in 2 then 4 copies, thus making 6 possible pairs, each gene being involved in 3 pairs. Depending on the method to identify ohnologs and their homologies, not all remaining pairs are identified with the same sensitivity and specificity. However, we show that despite their differences all 5 lists support the 2R hypothesis. Indeed, as for genes, each original chromosome is duplicated in 2 then 4 copies, and each chromosome of such a tetrad should possess 3 paralogous chromosomes. In *Amniota*, independently of the origin of the list used, a proportionality test (see methods) shows that CARs are, on average, linked by ohnolog pairs to 3 other CARS as expected (supplemental Figure S2). We thus decided to construct an improved consensus list of *Amniota* ohnolog pairs using all five lists and without favoring one in particular, under a constraint of compatibility with the 2R hypothesis.

A family of ohnologous *Amniota* genes is composed of genes that duplicated from a single pre-2R gene, and which remain in the *Amniota* genomes after potential deletions and segmental duplications that took place between *Vertebrata* and *Amniota*. In the simplest scenario, each gene in a family of ohnologs that was not subject to such deletions or additional duplications would be located on its own CAR, thus totaling 4 CARs per family. We call these genes linked by ohnologous relationships “networks of ohnologs”. Some networks may link fewer than 4 CARs, especially if one or two ohnologs were deleted post-2R. Some networks may connect more than 4 CARs, for example if a post-2R gene duplication relocated a fifth copy on a fifth chromosome. However we reasoned that the main reason for networks linking more than 4 CARs would be false positive ohnologs, i.e. genes that duplicated in another context than the 2R and were incorrectly included in an ohnolog list. In support of this, we noted that the intersection of all 5 lists, i.e. ohnologs independently identified by 3 different methods and thus considered the least prone to errors, never connect more than 4 CARs within a family. This list comprises 1,273 pairs of ohnologs (Figure 1A) grouped into 863 networks.

**Figure 1.**
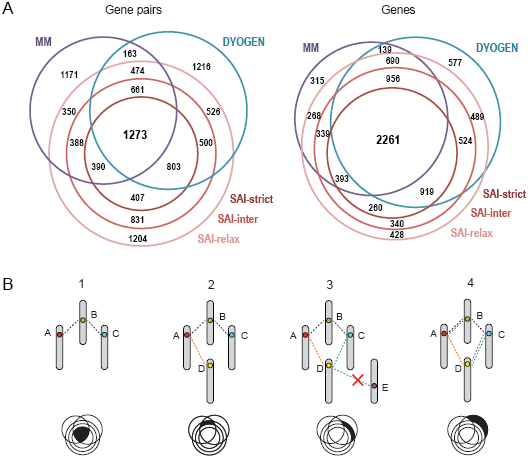
Identification of ohnolog pairs in the *Amniota* genome. (A) Comparison between five lists of ohnolog pairs in *Amniota*. Left: a Venn diagram of the sets of ohnolog pairs from five lists: the MM list (Makino and McLysaght 2010), the three SAI lists (Singh et al. 2015) and the DYOGEN list (this work). The numbers of pairs at the intersections of the lists are indicated. Right: a Venn diagram of the sets of ohnolog genes from the same lists as above. The overlap between the lists of ohnolog genes is higher than between the lists of pairs because the latter contain different pairs between the same genes. For example, two pairs A-B and B-C are in different lists (no overlap between lists) but gene B is common to both lists (1 gene overlap). The surface of the circles and their intersection are roughly proportional to the number of pairs of each list. (B) Schematic example of ohnolog pair selection. Step 1: from the initial list of 1273 gene pairs (black area in Venn diagram), a network of 2 pairs links 3 genes A, B and C, each on a different CAR. Step 2: pairs from a new sub-list are considered, a new gene pair B-D is added to the network. Gene D is on a fourth CAR. Step 3: A new list is considered, two new pairs are identified (D-C and D-E) but E is on a fifth CAR so pair D-E is discarded. Step 4: a new list is considered, two additional pairs supporting the network are identified.

We show further that this is a unique property of this intersecting list, because randomly drawing the same number of pairs from the complete union of lists (thus including potential false positives) always produces a list that has families connecting more than 4 CARs (Supplemental Figure S3). We call a network of ohnologs that does not connect more than 4 CARs a 2R-compatible network. Starting from the set of 1,273 high-confidence 2R-compatible pairs, we increased the list by adding pairs from sets of decreasing quality (Supplemental Table S1), as long as they either create a new network or they complement an existing network without connecting more than 4 CARs (Figure 1B).

The growing set thus remains 2R-compatible. This incremental process (Supplemental Table S1A) resulted in a list of 8,184 ohnologous genes, connected in 7,441 ohnolog pairs grouped as 2,973 networks, each in principle corresponding to one pre-2R gene (Supplemental Tables S2, S3, S4 respectively for the list of ohnolog genes along with their human descendants, the list of ohnolog pairs and the list of ohnolog networks).

We believe that this list of ohnologs is of high quality for a number of reasons. First, it is designed based on the gene content and synteny in the *Amniota* genome which is much closer to the 2R events, and thus where their signatures are read with greater accuracy, than on modern genomes which evolved for a further 326 million years. Second, it abides by a 2R-compatibility rule, i.e. no ohnologous gene family connects more than 4 CARs in its network. Third, the pairs are all phylogenetically consistent in that both genes in a pair always belong to the same Ensembl gene tree. This was the case by construction in the DYOGEN list, but not in the MM list (962 pairs of ohnologs, or 20%, belong to different trees) nor in the SAI lists (5%, 9% and 15% of pairs in the strict, intermediate and relaxed SAI lists respectively belong to different trees). Fourth, the two genes of a pair were allowed to be on the same CAR only if ≥ 90 genes apart to avoid spurious inclusions of tandemly duplicated genes. This criterion was also present by construction in the DYOGEN list but not in the MM and SAI lists (Supplemental Table S1B).

### Identification of post-2R ohnologous CARs

Using the improved list of ohnolog pairs, we wished to manually curate and re-assemble *Amniota* CARs into *Vertebrata* post-2R CARs, guided by the following reasoning. First, from *Vertebrata* to *Amniota*, chromosomes continued evolving during about 200 million years by fusions and fissions, thus respectively requiring breaks and merges of *Amniota* CARs to retrace these evolutionary steps. Second, the *Amniota* genome reconstruction by AGORA is neither entirely contiguous nor exempt from errors, also requiring respectively merges and breaks to improve the *Amniota* karyotype based on additional evidence.

With this is mind, we first built a dot-plot matrix of ohnologous genes in *Amniota* to identify CARs or segments of CARs (sub-CARs) that are strongly linked by ohnologs and thus likely to belong to duplicated *Vertebrata* chromosomes (Supplemental Figure S4). Areas of higher density occur between specific pairs of CARs, readily identifying ohnologous regions of the *Amniota* genome. To quantify this observation, a proportion test (methods) assigned a p-value to each pair of CARs, reflecting how unlikely their number of shared pair of ohnologous genes is, compared to a random distribution.

Analysis of the dot matrix shows that five CARs (5, 118, 40, 97, 46) can be divided in two segments: each possesses ohnologs with a distinct set of CARs (e.g. CARs 5, 40, 118 in supplemental Figure S4). Such situations are expected if fusions of chromosomes took place since *Vertebrata* (CARs 5, 118; Supplemental information) or if AGORA incorrectly assembled two disjoint *Amniota* chromosomes (CAR 40, 97, 46; Supplemental information). We therefore split each of the five CARs in two CARs according to their ohnolog distribution. After this step, the assembly of *Amniota* contained 59 CARs of at least 50 genes.

We next set out to identify sets of duplicate CARs that arose from one pre-2R chromosome during the two WGD. After establishing pairwise links between CARs with the above proportionality test, we built a graph of CARs connected by ohnologous genes with a Bonferroni adjusted p-value threshold of 5.10^−2^. Ideally, the graph should correspond to disjoint tetrads, i.e. independent sets of four CARs all ohnologous pairwise. However, this may not be so for at least 2 reasons: (1) one CAR of the tetrad may be absent (not retained, or not assembled in the *Amniota* reconstruction, or not found because of biased massive loss of ohnologs) and (2) evolutionary events such as fissions, fusions and rearrangements are likely to have occurred between the two rounds of WGD (Nakatani et al., 2007; Putnam et al., 2008), making the final pattern of links more complex than a simple tetrad. To account for this complexity, we conservatively nucleated our search starting from triads (3 CARS connected pairwise) instead of tetrads. Limiting the graph to the 59 CARs of at least 50 genes linked by ohnologs, we identified 8 disjoint sets of CARs containing a minimal pattern of triad. With the same p-value threshold, we next added smaller CARs of at least 5 genes to these sub-graphs in order to complete triads into tetrads. Most sub-graphs contained ohnolog links suggesting that some CARs should be assembled as part of the same *Amniota* chromosomes: they belong to the same cluster, they share links to common CARs and are not linked to each other. In addition, their assembly is structurally supported by at least one of three additional evidences (Table 1, supplemental information). This observation led to 23 assemblies of two to five CARs within all but one cluster (Table 1; supplemental Figures S7 to S14 B).

**Table 1:** Assemblies of *Amniota* CARs within clusters. Each line corresponds to one assembly of two to five CARs (column 1) within one cluster (column 2). Columns 3 to 5 indicate which structural conditions are fulfilled. The last three columns indicate additional but dispensable information to support the assembly. “v.84” refers to Ensembl version 84, gar-chicken 1:1 indicates that scaffolds map to homologous gar-chicken chromosomes; DCS, Double Conserved Synteny with Medaka. A full description of the table can be found in supplemental information

| Groups of assembled CARs | Cluster | Physical link (one or more required) |  |  | Additional syntenic observations |  |  |
| --- | --- | --- | --- | --- | --- | --- | --- |
|  |  | AGORA scaffolding | Same CAR in v.84 | Chicken (+) or human (x) chromosome | Same gar Linkage Group | gar-chicken 1:1 | DCS/ Medaka |
| 6, 39, 140 | #1 | + | - | + | + |  | + |
| 80, 120 | #2 | + | + | + | + |  | + |
| 107, 207 | #2 | + | - | + | + | + | + |
| 5a, 152 | #2 | - | - | + | + |  | + |
| 254, 48 | #2 | - | - | x | n.d. |  | + |
| 65, 53, 103 | #3 | + | - | + | + |  | + |
| 114, 195 | #3 | + | + | + | + | + | + |
| 26, 102 | #3 | + | + | + | + | + | + |
| 50, 85, 84 | #3 | - | + | x | + |  | + |
| 67, 111, 83 | #3 | 67 & 83 | - | + | + | + | + |
| 146, 450, 456, 393, 52 | #3 | - | 146 & 450 | x | n.d. |  | 146, 393 & 52 |
| 117, 137, 256 | #4 | 117 & 137 | 137 & 256 | + | + | + | + |
| 204, 75, 55 | #5 | 204 & 75 | - | + | + |  | + |
| 129, 261, 97a | #5 | 261 & 97 | - | + | + |  | + |
| 59, 420 | #5 |  | - | x | + |  | + |
| 3, 22 | #6 | + | + | + | + | + | + |
| 60, 148 | #6 | + | + | + | + |  | + |
| 10, 240, 2 | #6 | 10 & 240 | - | + | + |  | + |
| 5b, 40b | #6 | - | + | + | + |  | + |
| 118a, 192, 14, 172 | #7 | 14 & 192 | - | + | + |  | + |
| 66, 46a | #7 | - | - | x | + |  | + |
| 123, 46b | #7 | + | + | + | + |  | + |
| 441, 430, 93 | #7 | - | + | x | n.d. |  | + |

For example, cluster #4 initially consisted of 5 CARs larger than 50 genes, of which 3 were linked in a triad (Supplemental Figure S10A). The tetrad was completed with the addition of CAR 82 (9 genes), which was linked to 4 of the 5 initial CARs with significant p-values (Supplemental Table S5). Then, of the 5 initial CARs, 3 fulfilled the conditions for being assembled in a single larger CARs (CARs 117, 137, 256): they belonged to the same cluster #4, shared links to CARs in common and did not show any links to each other, leading to a perfect tetrad (Supplemental Figure S10B). In addition, AGORA scaffolded CARs 117 and 137 together, fused CARs 137 and 256 together when using a more recent version of Ensembl, the three CARs map to the same chicken and gar chromosomes and to the same medaka pair of chromosomes linked by double conserved synteny (Supplemental Table S5), altogether strongly supporting this assembly. In fact, all assembled CARs project on the same gar (*Lepisosteus oculatus*) linkage groups (LG) (Braasch et al. 2016), except for three sets of small CARs for which the gar LG could not be identified (Table 1). All clusters could be resolved into one to four tetrads (Figure 2A and supplemental Figures S7 B to S14 B). The resolution of CARs in tetrads is much clearer with the *Amniota* karyotype than what could have been achieved using the human genome (Figure 2B and 2C). The proportionality test links each *Amniota* CAR on average to 3 other CARs almost independently of any p-value threshold, whereas human chromosomes are much more sensitive to the p-value threshold. Indeed in the human genome, the expected average of three partners per chromosome is reached at a p-value of 1.10^−09^, a stringent threshold that leaves 7 human chromosomes unlinked to any other chromosome. The proportionality test also converges toward three partners faster on the assembled *Amniota* genome than in the non-assembled genome (respectively Figure 2C and supplemental Figure S2F).

**Figure 2.**
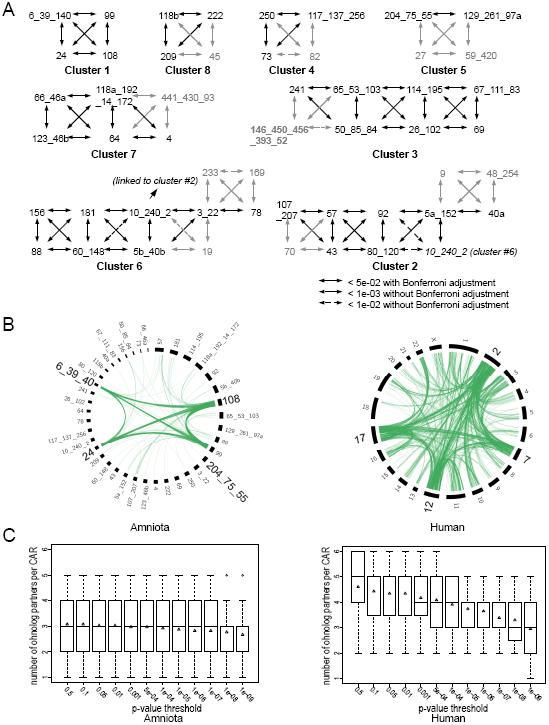
Assembly of the Amniota karyotype in tetrads (A) Eight clusters of CARs in tetrads. CARs are numbered arbitrarily and are joined by underscores in an arbitrary order when assembled. The letters ‘a’ or ‘b’ indicates that the CAR has been split in two segments. Double-headed arrows represent the ohnology links Solid lines are links supported by a Bonferroni-adjusted p-value of less than 0.05 (proportion test), dotted lines are links that are supported without the Bonferroni adjustment. Numbers in black indicate CARs of at least 50 genes, while smaller CARs (< 50 genes) are in grey. (B) Circos plot (Krzywinski et al. 2009) showing the pairs of ohnologs involving each of the four chromosomes (*Homo sapiens*) or CARs (*Amniota*) of the tetrad carrying the Hox genes (Cluster 1 in A). The pairs of ohnologs in the human genome were the descendants of those of *Amniota* (6,121 pairs of human ohnologs vs. 7,441 pairs of amniote ohnologs). The human *Hox* cluster tetrad is mainly composed of human chromosomes 2, 7, 12 & 17. The *Amniota Hox* cluster tetrad is composed of CARs 108, 24, 99 and 6_39_140. An ohnolog pair is represented (green lines) between two *Amniota* CARs or two human chromosomes only if at least one of the two genes of the pair falls on a chromosome/CAR of the tetrad. The *Amniota Hox* CARs are involved in 634 pairs, while the human *Hox* chromosomes are involved in 2,171 pairs of ohnologs. This figure shows that the reconstruction of *Amniota* ancestor displays a clearer picture of the 2R than the human genome. (C) Ohnolog partners per chromosome/CAR in the human (left) and *Amniota* (right) genomes. Each boxplot shows the distribution of the number of CARs (*Amniota*) or chromosomes (Human) found to be ohnologous to a given chromosome/CAR by the proportionality test. The x-axis shows the Bonferroni adjusted p-value thresholds used to select ohnologous chromosome/CARs. Triangles indicate the average number of partners. The *Amniota* genome shows a clear and stable distribution of three partners per CAR across a wide range of p-values, as expected after two WGDs where chromosomes are linked in tetrads. In contrast, the distribution in *Homo sapiens* shows that extremely low p-value thresholds must be used to reach the expected average of three partners, justifying the fragmentation of the human genome as described in (Nakatani et al. 2007).

### Chromosome evolution during two whole genome duplications

A one-tetrad cluster implies a single evolutionary scenario without large chromosomal rearrangement between the two WGD (Figure 3A). Two adjacent tetrads however, can be explained by one of two scenarios, each with the same degree of parsimony: a fission of two CARS or the fusion of two CARs at some point between the 2R (Figure 3B).

**Figure 3.**
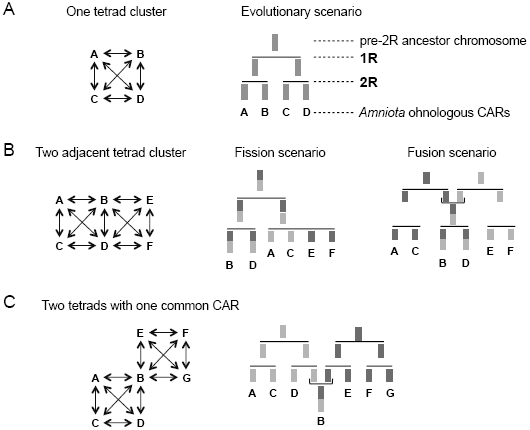
Evolutionary scenario models (A) A single evolutionary scenario explains a one-tetrad cluster of ohnologous CARs. (B) Two equally possible evolutionary scenarios can explain a cluster of ohnologous CARs comprising two adjacent tetrads: a fission or a fusion of chromosomes could have occurred between the 2R. In each case, the B and D chromosomes each possess two distinct parts (dark and light grey) homologous to distinct chromosome sets. B and D are therefore common to two tetrads (C) A chromosome fusion after the two WGDs explains clusters with two tetrads joined via a single CAR.

Without additional information, the choice between the two scenarios is not possible and yet has important consequences because in the case of a fission, it implies a single pre-2R chromosome while in the case of a fusion it implies two pre-2R chromosomes (Nakatani et al. 2007). As previously observed (Jaillon et al. 2004; Kellis et al. 2004), a non-duplicated outgroup species would be helpful to discriminate between the two possibilities. Amphioxus (*Branchiostoma floridae*) is a cephalochordate with a slowly evolving genome (Putnam et al. 2008; Louis et al. 2012), which represents such a suitable outgroup. Evidence for a fission scenario would be provided if genes from a single amphioxus scaffold had orthologs distributed over a single reconstructed pre-2R chromosome. A direct comparison between the *Amniota* and the amphioxus genomes failed to support such fission scenarios, in any of the adjacent tetrads (data not shown). The absence of signal may however be caused by the fragmented nature of the amphioxus genome assembly (N50= 138 genes). To circumvent this problem, we used 17 reconstructed chordate linkage groups (CLG) that have been proposed to each represent the unordered gene content of an ancestral chordate chromosome (Putnam et al. 2008). This reconstructed proto-karyotype precedes the 2R events by less than 50 million years and is located at a much shorter evolutionary distance to *Vertebrata* than the extant amphioxus genome.

Under the same rationale as for amphioxus scaffolds, we compared the gene content (19,020 human genes) of the 17 CLGs to the human genes descended from the genes content of *Amniota* tetrads (see below). Here again, we could not find evidence for chromosome fissions occurring between the 2R: remarkably, each CLG was associated to one predominant tetrad. However we found evidence for a fusion between CLG 6 and CLG 7 before the 2R, because tetrad #6a corresponds to both CLGs (Supplemental Figure S16, and supplemental Table S7). We can therefore conclude that the pre-2R karyotype comprised 17 chromosomes, giving rise to chromosomes after the first WGD, followed by 7 fusions (27 chromosomes), which were duplicated again in the second WGD leading to 54 Vertebrata chromosomes, at the origin the approximately 60,000 species of vertebrates. It was immediately followed by four fusions leading to 50 chromosomes at the origin of *Euteleostomi*. A fifth fusion occurred specifically in the *Amniota* lineage, leading to 49 ancestral chromosomes in the *Amniota* genome (Figure 4; Supplemental information).

**Figure 4.**
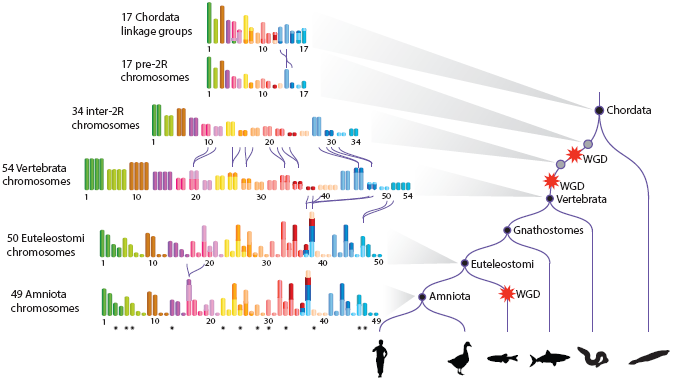
Reconstructed evolutionary history of karyotypes from *Chordata* to *Amniota*. On the right, a simplified species tree of the *Chordata* is shown, with WGD events depicted by red stars. The five lineages represented from left to right are mammals, birds, teleost fish, cartilaginous fish, cyclostomes (lamprey, hagfish), cephalochordates (amphioxus). On the left, successive reconstructed karyotypes are shown, with one color for each of the 17 pre-2R chromosomes. The length of each pre-2R chromosome is proportional to its number of genes. For the 17 Chordate Linkage Groups (CLGs) of (Putnam et al. 2008), the size of the colored segment is proportional to the number of genes that are found in the intersection of the CLG with a pre-2R chromosome, although segments corresponding to < 10% of the number of genes of the CLG were omitted for clarity (supplemental Table S7). The inter-2R karyotype was deduced from the pre-2R karyotype and the seven chromosome fusions are shown with black curvy lines joining the fused chromosomes. The *Euteleostomi* karyotype was deduced from the Vertebrata karyotype after four chromosome fusions (supplemental information). The lengths of the *Euteleostomi* chromosomes are proportional to the number of genes in the homologous *Amniota* CARs. Finally the *Amniota* karyotype differs from that of *Euteleostomi* by only one chromosome fusion. The *Amniota* chromosomes were numbered from 1 to 49 (Supplemental Table S11 for correspondence with the CARs and number of genes). Black stars under 12 Amniota chromosomes denote predicted ancestral microchromosomes.

In order to complete the pre-2R vertebrate ancestor genome reconstruction, we assigned sets of ancestral genes to each of the 17 predicted chromosomes. The AGORA algorithm reconstructed a total of 10,093 ancestral genes for the *Olfactores* ancestor, which is the common ancestor of vertebrates and urochordates and the closest in our dataset to the reconstructed pre-2R genome. We applied a conservative procedure (methods) to assign 5,043 of these *Olfactores* genes (50% of the 10,093 ancestral *Olfactores* genes) to the 17 pre-2R chromosomes (supplemental Table S12). This set of ancestral genes provides a direct connection to the human genome, through their 8,378 human descendent genes (Figure 5). The structure of the 17 pre-2R chromosomes is still strikingly apparent, with some human chromosomes almost entirely composed of genes from a single pre-2R chromosome (eg. chromosomes 14 and 15). We measured the degree of conservation of the post-2R ohnolog content in scanning windows across human chromosomes. Three regions overlapping the Hox clusters A, B and D stand out (and Hox C to a lesser degree; Figure 5), in line with the known functional importance associated with the clustering of these ohnologs (Krumlauf 1994).

**Figure 5.**
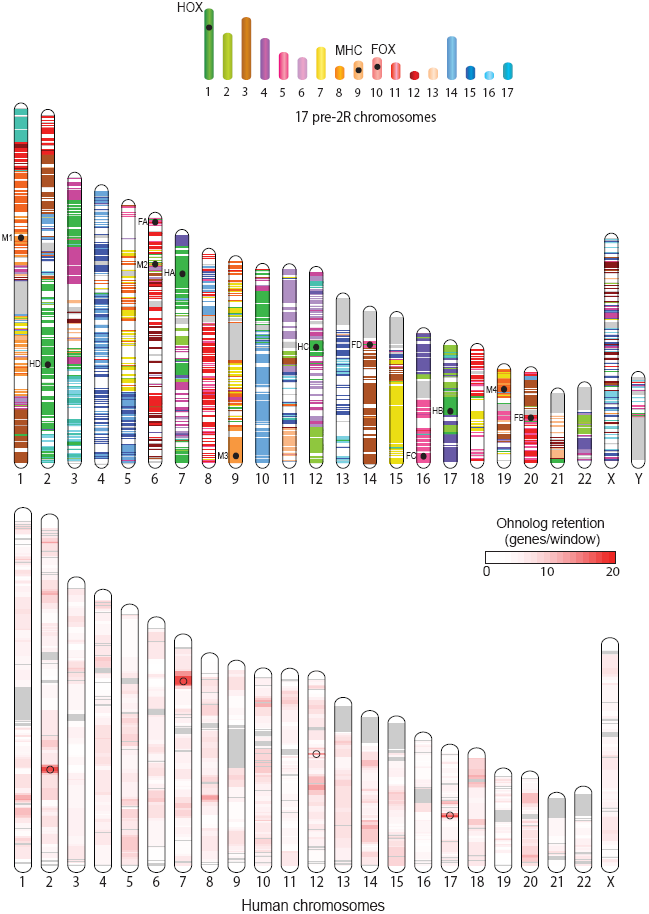
Comparison between the pre-2R karyotype (top) composed of 17 chromosomes and the human karyotype (middle). The 8282 known human descendent genes of pre-2R genes are drawn at their position in the human genome with the colour of their pre-2R ancestral chromosome. The position of 12 extant clusters (4 HOX, 4 FOX and 4 MHC) descending from a single clusters in pre-2R chromosomes are indicated by a black circle and a 2-character identifier (M1, M2, M3, M4 for MHC clusters, F1, F2, F3, F4 for FOX clusters, HA, HB, HC, HD for HOX clusters). A second human karyotype (bottom) shows, in a white-to-red scale, the number of ohnologs in scanning windows of 50 genes in steps of 10 genes. Open circles denote the position of HOX clusters. Human chromosomes are drawn to scale, in Mb. Pre-2R chromosomes are drawn as in Figure 4, in proportion to the number of genes assigned to each. The order of genes in pre-2R chromosomes being unknown, the positions of the 3 pre-2R gene clusters are arbitrary.

The four clusters originate from pre-2R chromosome 1, and we examined in the same light other paralogous clusters that have been proposed to originate from a single pre-2R locus. The MHC region on human chromosome 6 contains a number of genes unrelated to immune functions but which possess ohnologs in 3 other loci on chromosomes 1, 9 and 19 (Danchin and Pontarotti 2004). All 4 regions descent from pre-2R chromosome 9. Similarly, the loci containing FOX gene clusters have been compared and found to share paralogs suggestive of en-bloc duplications early in vertebrate evolution (Wotton and Shimeld 2006). We confirm here their unique origin on pre-2R chromosome 10. In contrast, no common origin was found for the different cluster of imprinted genes in the human genome (Morison and Reeve 1998; Dunzinger et al. 2007), in line with their progressive appeareance later in therian mammals (Renfree et al. 2009).

## DISCUSSION

We analyzed 61 animal genomes to reconstruct the evolutionary history of genes and chromosomes at the time when vertebrate were about to diversify into more than 60,000 species. In contrast to analysing extant genomes as in previous studies, we first reconstructed the *Amniota* genome to a high degree of continuity, as a stepping-stone to identify the signature of chromosome duplications more clearly. Our rationale is that *Amniota* should be devoid of the noise caused by the numerous rearrangements that took place during the following 326 million years of evolution. Indeed, the benefit of this approach can be seen when comparing the distribution of ohnologs in the *Amniota* genome versus their descendants in the human genome, for example in the chromosomes carrying the *Hox* clusters (Figure 2B). The expected four-way association between duplicated chromosomes is striking in *Amniota* but blurred in the human genome. Statistical confidence and power are thus higher, enabling a higher resolution of chromosome events than was previously possible. In turn, this property of the data enables a straightforward reconstruction strategy of pre-duplication genomes, without requirements for complex statistical steps or algorithmic developments (reviewed in (Muffato and Roest Crollius 2008; Nakatani and McLysaght 2017)).

Progress was made on understanding early vertebrate genome evolution when (Nakatani et al. 2007) reconstructed a vertebrate pre-duplication ancestral genome by segmenting the human genome into conserved vertebrate linkage groups. These were compared with other vertebrate genomes, especially with medaka, using tunicate and sea urchin genes to define ohnologs. The authors inferred a karyotype of the pre-2R vertebrate ancestor, but could not resolve inter-WGD chromosome fusions or fissions, leaving the number of pre-2R chromosomes to a range comprised between 10 and 13 chromosomes. Here, using the *Amniota* instead of the human genome, we reconstruct 17 chromosomes for the pre-2R genome and find evidence for 7 fusions between the 2R but none for fusions. We identify new chromosomes and find major differences between the two karyotypes (supplemental Table S8). Our study provide an improved picture of ancestral vertebrate genome evolution in part because it is based on the reconstructed *Amniota* genome, and relied on information from more species and more recent genome annotations. For example the gar genome was more informative than the synteny with the medaka genome (Kasahara et al. 2007) because it is exempt of the numerous chromosome fusions that occurred just before the teleost WGD (Braasch et al. 2016).

The same authors (Nakatani et al. 2007) also reconstructed an *Osteichthyes* post-2R genome corresponding here to the *Euteleostomi* ancestor, using a 2-of-3 rule relying on conservation between two of three genomes: the teleost pre-WGD, the chicken and the vertebrate pre-2R genome. Here however our *Euteleostomi* ancestor is substantially different from this earlier study, since we describe an *Euteleostomi* genome of 50 chromosomes, not 31. This difference mainly comes from fusions that we believe were incorrectly inferred from chicken-teleost genome comparisons, which were confounded by high rates of chromosome fusions in the lineage leading to the ancestral teleost (Braasch et al. 2016). Given that the ancestral teleost fish possessed only 13 chromosome, the 50 chromosomes inferred here in the ancestral Euteleostomi suggest that the rate of chromosomes fusions in the lineage leading to teleosts must have been more intense than previously thought, of the order of 37 fusions in 100-150 million years. Similarly, the ancestral eutherian karyotype probably consisted of 23 pairs (Ferguson-Smith and Trifonov 2007), suggesting a consistent pattern of karyotype reduction by chromosome fusion after the 2R whole genome duplications.

We examined several questions using the improved picture of ancestral vertebrate chromosome described here. First, it should be noted that neither the reconstruction of the *Amniota* genome nor the establishment of ohnolog pairs described here make any assumption about the existence of two successive WGDs early in vertebrate evolution. Indeed, the criteria used to select ohnologs only rely on duplication dates and on local synteny, leaving open the possibility that segmental duplications, one single WGD or two WGDs have occurred during early vertebrate genome evolution. The fact that *Amniota* CARs readily associate to form tetrads when using duplicated genes as links (Figures 2), is a striking confirmation of the 2R scenario. This scenario is in fact largely consensual today, but the debate was recently re-opened (Smith and Keinath 2015) after a reconstruction of the pre-2R vertebrate genome using the lamprey (*Petromyzon marinus*) genome sequence (Smith et al. 2013) and the chicken genome. Lamprey is an Agnathan (jawless vertebrate), a sister group to Gnathostomes (Heimberg et al. 2010) to which Amniote belong, and both groups share the pre-2R genome as common ancestor (Figure 4). The lamprey genome is composed of almost 100 chromosomes and its lineage has separated from the Gnathostomes soon after the 2R. Redundant gene copies from the 2R have therefore likely been lost independently in both lineages, leaving the possibility to find homologous chromosomes between the lamprey genome and a Gnathostome (here chicken) genome as long as both sets of chromosomes descend from the same pre-2R chromosome (Smith and Keinath 2015). The result suggested that the ratio of ancestral pre-2R chromosomes to chicken chromosomes was mostly 1:2 and less frequently 1:4 or even 1:3, thus supporting a single WGD combined with additional segmental duplications. However, when replicating the above study with our reconstructed *Amniota* genome instead of chicken, a clear majority of 1:4 patterns appears (Supplemental Figure S17, supplemental information), hence confirming the occurrence of 2 successive WGDs. This further emphasizes the benefit of using ancestral genome reconstructions as intermediates when investigating such ancient evolutionary events (about 450 millions years before present).

The reconstructed *Amniota* genome is not complete, in that the 49 chromosomes only contain 80% of the 15,854 *Amniota* genes that belong to CARs (of 2 genes and more), and 3932 genes remain as singletons, i.e. ancestral genes for which no adjacency could be identified. We note that although all chromosome tetrads corresponding to pre-2R chromosomes are complete, the 49 *Amniota* chromosomes display large differences in gene numbers: the largest contain 862 genes (chromosome 37) and the smallest only 16 genes (chromosome 49). This could reflect either a more intense process of gene inactivation and loss on chromosomes with fewer genes, or a more intense rate of rearrangement on those chromosomes, leading to greater difficulties in reconstructing them. Small chromosomes with few genes do not follow any noticeable pattern in their distribution among tetrads, which could have otherwise indicated systematic biases in gene deletion during rediploidisation. We further examined if chromosomes could be paired within a tetrad, as expected if gene loss following the first WGD left a distinct pattern on the two ohnologous chromosomes, that would have propagated to the two duplicates resulting from the second WGD. Here again such a pattern is not noticeable, which may indicate that the two WGDs took place in rapid succession, as suggested before (Furlong and Holland 2002), leaving little time for gene deletions (diploidisation) to leave their imprint. More high quality genome sequences from extant amniotes are required to improve the reconstruction of their ancestral genome and identify more subtle patterns left by the 2R. However, the current Amniota reconstruction and its ohnolog gene annotation provides a foundation for new studies that should help resolve important questions, including complex phylogenetic histories (Lynch and Wagner 2009).

An interesting question was raised during the analysis of the gar genome (Braasch et al. 2016), when authors noticed a frequent 1:1 relationship between gar and chicken chromosomes, including micro-chromosomes. Micro-chromosomes are unusual because of their small sizes (usually below 20 Mb in chicken), their high GC content, high recombination density and high gene density. The gar-chicken comparison suggests that micro-chromosomes are ancestral features in *Euteleostomi*, which in turn raises the question of their origin through the 2R duplications. Twelve gar and chicken micro-chromosomes are homologous and can parsimoniously be considered ancestral to *Euteleostomi* (Supplemental Table S10). Their distribution in the tetrads resulting from the 2R does not follow a noticeable pattern, i.e. they are distributed among all tetrads more or less randomly (Figure 4). We can conclude that micro-chromosomes do not originate from a set of pre-2R micro-chromosomes, but that this characteristic started evolving at least after the first WGD.

The pre-2R karyotype and its evolution described here provide a framework to study the impact of the 2R in extant vertebrate genomes, especially via the set of phylogenetically consistent ohnolog families that underlie the analysis. The pre-2R karyotype also provides a reference to study the diversification of vertebrate genomes from the standpoint of their common ancestor.

## METHODS

### Reconstruction of the *Amniota* ancestral genome

The *Amniota* genome was reconstructed using the AGORA algorithm (Berthelot et al. 2015) (https://github.com/DyogenIBENS), which is routinely used to reconstruct ancestral gene order presented in Genomicus since Ensembl release 53. The Amniota reconstruction used here is available for download on the ftp site of the Genomicus webserver (Nguyen et al. 2017) (ftp://ftp.biologie.ens.fr/pub/dyogen/genomicus/69.01/). Briefly, AGORA used 61 extant metazoan genomes (gene order, gene orientation, gene trees) from release 69 of Ensembl (2012/11/15) to build an adjacency graph where ancestral genes are vertices and ancestral adjacencies are edges weighted by the frequency of their conservation. The graph is then linearized along the edges of maximal weight to produce Contiguous Ancestral Regions (CARs). An AGORA reconstruction of the *Amniota* genome based on Ensembl version 84 (2016/03/15) that included the *Lepisosteus oculatus* (spotted gar) genome (Braasch et al. 2016) was also used for comparison. The species tree we referred to can be found at:
http://www.genomicus.biologie.ens.fr/genomicus-69.01/data/SpeciesTree.pdf

### Identification of ohnolog gene pairs in *Amniota*

Human ohnolog gene pairs from (Makino and McLysaght 2010) based on Ensembl 52 data were assigned to their Ensembl version 69 gene trees using Ensembl gene IDs. Their *Chordata* ascendants were used to build the MM list of ohnolog gene pairs. Similarly, we downloaded three lists of ohnolog gene pairs from http://ohnologs.curie.fr/ described in (Singh et al. 2015), each list corresponding to a different degree of confidence level. A non-redundant list of *Amniota* ascendant genes was identified in Ensembl version 69 by sequentially identifying the *Amniota* ascendant genes of the ohnologs in the six extant amniote genomes (data downloaded on October, 27, 2014). Finally, we built the DYOGEN list by identifying ohnolog pairs directly in the *Amniota* reconstructed genome starting from ancestral genes that were duplicated between the *Euteleostomi* ancestors: these candidate pair were considered ohnologs if another candidate pair from a different gene tree could be found on the same pair of CARs at a distance ≤ N genes (to account for massive duplicate loss after a WGD): the parameter N was made vary to optimize both sensitivity and specificity and fixed to 45 genes. A pair of ohnologs was allowed to occur between genes of the same CAR if they were located at a distance of 2N genes (90 genes, to account for possible rearrangements on the *Amniota* lineage), which added only 44 pairs. Next, an integrated, high quality list of ohnolog gene pairs was built from these 5 lists. Starting from their intersection consisting of 1273 pairs, we built disjoint graphs of ohnologs connected by ohnologous relationships. Replacing genes by their CARs, these graphs never involved more than four CARs of at least ten genes. To this core list, we sequentially added lists of pairs present in several or only one of the five initial lists, removing at each step the newly added pairs that would build ohnolog networks involving more than four CARs (Supplemental Table 1). The order in which these added lists were considered was established using the levels of confidence of the SAI lists, the number of lists where the pairs were found and the 2R-compatibility criterion: a list was of better quality if it created fewer ohnolog networks of more than four CARs (per added pair) when added to the current validated list. Two properties of the DYOGEN list were maintained along the process: (1) the two ohnologs of a pair were allowed to be located on the same CAR only if they were ≥ 90 genes away from each other; (2) the two ohnologs of the pair had to belong to the same Ensembl gene tree.

### Identification of ohnolog CARs in *Amniota*

Given the distribution of ohnologs on the *Amniota* CARs, a proportion test was performed (prop.test function in R) between each pair of CARs to estimate if the corresponding CARs shared more ohnolog pairs than expected by chance. Bonferroni adjusted p-values of 0.05 or less were considered significant. However p-values obtained without the Bonferroni correction were also examined if CARs were included in a tetrad with a least one significant Bonferroni adjusted p-value.

### Attribution of *Olfactores* genes to the 17 pre-2R chromosomes

Gene trees from version 69 of Ensembl were analyzed to identify 10,093 nodes (genes) in the *Olfactores* genome, which were assigned to the 17 pre-2R tetrads according to the distribution of their descendent *Amniota* genes in the *Amniota* CARs used to build ohnologous tetrads. In the simple case of one-tetrad clusters, all *Olfactores* genes with descendants exclusively in the 4 CARs of a tetrad were included in the corresponding pre-2R chromosome. *Olfactores* genes with descendants in more than one tetrad were considered ambiguous and excluded. In the more complex cases of multi-tetrad clusters, the same principle was applied but restricted to ohnolog pairs when CARs belonged to two adjacent tetrads. In order to maximize the number of ancestral genes in the pre-2R chromosomes, we used all possible small CARs (≤ 50 genes) that could be assembled to the CARs of the tetrads, respecting all criteria of the 23 previous assemblies (Supplemental Table S6)

### Human genes derived from ancestral chordate linkage groups (CLGs)

The coordinates of the 120 human chromosome segments from Tables S1 and S14 in (Putnam et al. 2008) were updated from the hg18 to hg19 version of the human genome using the UCSC *liftOver* utility. Twenty-two segments were not converted by *liftOver* and were further fragmented in 100 sub-segments of equal size, which were mapped again to hg19 in order to recover as many genes as possible (supplemental Figure S15).

## ACKNOWLEDGMENTS

We wish to thank Pierre Vincens for the coordination of computing resources and Lucas Tittmann for assistance with statistics. This work was supported by grants from the French Government and implemented by ANR (ANR-10-BINF-01-03 Ancestrome, ANR– 10–LABX–54 MEMOLIFE and ANR–10–IDEX–0001–02 PSL* Research University).

